# Understanding the Role of Toggle Genes in Chronic Lymphocytic Leukemia Proliferation

**DOI:** 10.1101/2025.03.31.646494

**Authors:** Olga Sirbu, Gunjan Agarwal, Alessandro Giuliani, Kumar Selvarajoo

## Abstract

Cancer cell populations, such as chronic lymphocytic leukemia (CLL), are characterized by aberrant proliferation and plasticity: cells may switch between states so increasing population heterogeneity. Previous works have shown that gene expression noise can impact cell-state transition. To gain better insights into transcriptome-wide expression dynamics and the effect of noise on state transition, here we investigate RNA-Seq data of proliferative (PC) and non-proliferative (NPC) CLL cells. Various data analytics were applied to the whole transcriptome, switch-like toggle (ON/OFF) genes, temporal differentially expression (DE) genes, and randomly selected genes. Collectively, we identified 2713 temporal DE genes (DESeq2 with 4-fold, *p* < 0.05) and 1704 toggle genes shaping the differentiation process over a period of 96h, with 604 overlapping genes between them. Despite their lower numbers compared to DE, toggle genes contributed significantly to gene expression noise in both cell types. Toggle gene analyses revealed the enrichment of genes involved in processes such as G-alpha signaling and muscle contraction as proliferation related RHO-GTPase, interleukin and chemokine signaling, and lymphoid cell communication. Thus, toggle genes, although being random (at single cell level) ON/OFF genes, can show (at population level) a symmetry breaking favoring one of the two ON/OFF states so contributing to gene expression functional variability. These results suggest that toggle genes play an important role in shaping cell population plasticity.

## Introduction

Chronic lymphocytic leukemia (CLL) is the most common type of leukemia in adults, with a median age of diagnosis and onset of 70 years^1^. It is characterized by the uncontrolled proliferation of monoclonal lymphoid cells, specifically transformed mature CD5+ and CD23+ lymphocytes which are impaired in their function^2,3^. Due to the heterogeneous nature of CLL, current treatment approaches for the disease are complex and suboptimal^2,4–8^. Previously, it has been observed that tumors can leverage genetic, epigenetic, and stochastic variability to foster the necessary plasticity that leads to resistance and treatment evasion^9–12^. While CLL is known to exhibit significant clonal and metabolic plasticity, its transcriptomic plasticity remains underexplored. Thus, transcriptome-wide analytics, that are capable of tracking systemic responses in gene expression, is necessary and it offers an important avenue for the study of CLL plasticity.

The construction of gene expression landscapes ^13,14^ allows to understand transcriptome-wide expression dynamics, especially in the context of cancer. This approach implies the conceptualization of living cells as dynamic systems that occupy specific states at any given moment. As cells undergo dynamic processes they move through the landscape, eventually tending towards conditions of stability or equilibrium, known as “attractors” (Figure 1A)^8,13,15–17^. Thus, the gene expression trajectories that cells follow as they move through the expression landscape are important for cell-fate decision making.

**Fig. 1.**
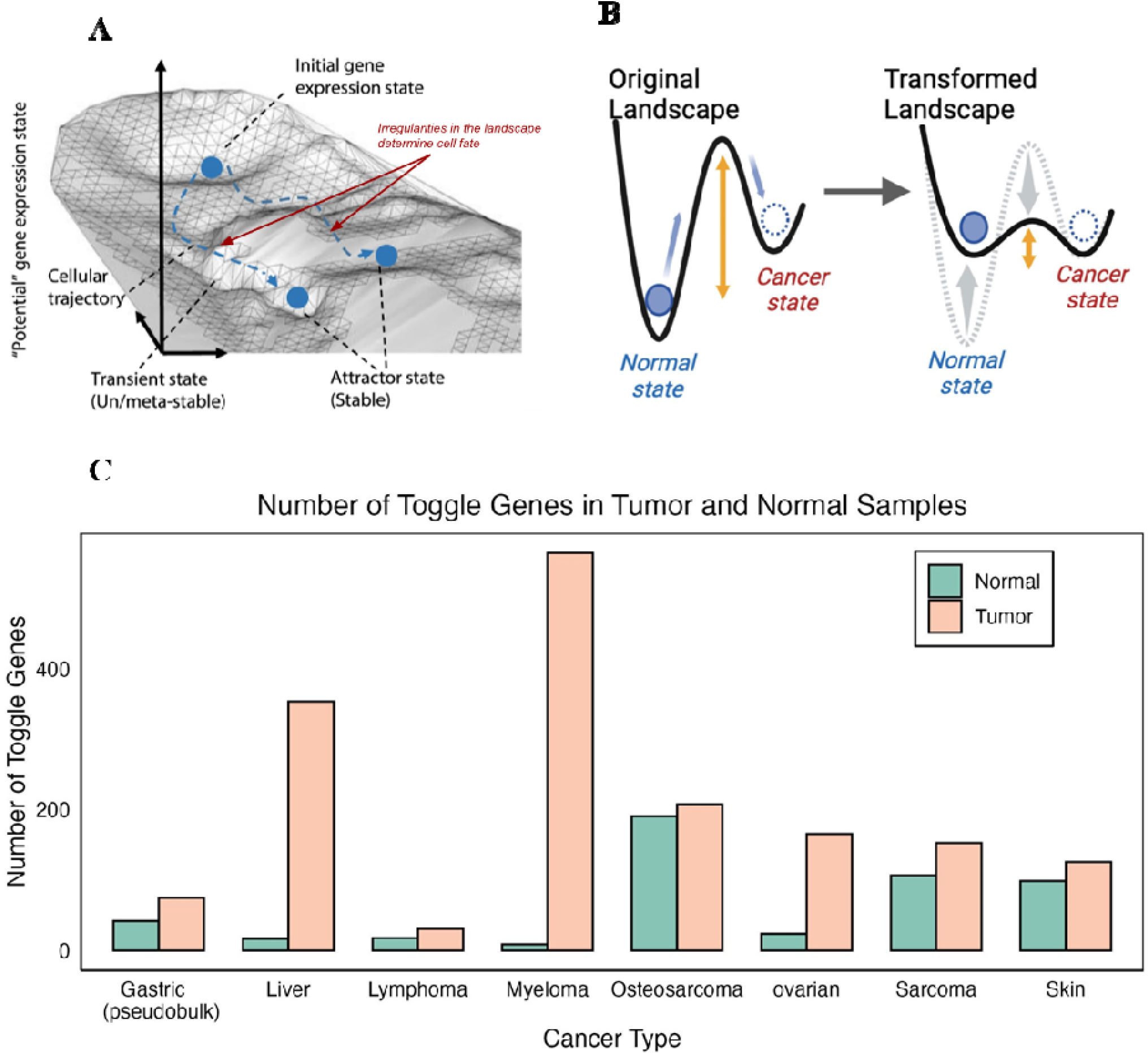
(A) Transcriptome expression landscape shows how cells follow certain trajectories to settle into different attractor states, figure adapted from ^16^. (B) Normal (left) and transformed (right) simplified cell fate landscape which shows that cells require larger perturbations (yellow arrows) to exit their current normal state and have the potential to fall into the cancer attractor. In a transformed landscape, the energy barrier required changes due to changes in attractor depth and thus state changes are more likely to occur. (C) Breakdown of toggle genes extracte from normal (orange) and tumor (green) samples shows a higher incidence of toggle genes in tumor samples across 8 investigated cancer sets (Table S2). Only datasets containing paired normal and tumor samples were selected t allow for direct comparison, however, the incidence of toggle genes in cancer is ubiquitous.

For cancer, we can think of a simplified cell-fate landscape with only two attractors: a normal state, and a cancer state. Under normal circumstances, cells are more likely to settle into the normal cell attractor, and very large perturbations are necessary to cause a cell to move to the cancer attractor (Figure 1B, left). However, cancer cell transcriptomes exhibit a level of plasticity that endows them with unpredictable behaviors and patterns, rarely seen in normal healthy cells^17–20^. In the case of an altered landscape (being this alteration coming from diverse initial causes), the perturbation required to exit the normal attractor and settle into a new cancer state is significantly smaller (Figure 1B, right). Therefore, external perturbations such as gene expression noise, can play major roles in shaping cancer states^4,18,21–24^.

Previous works have shown that gene expression noise plays a significant role in producing diversity and shaping complex biological processes^21,25,26^. During cell fate decision making, transcriptome-wide noise has been associated with controlling lineage choices in mammalian progenitor cells, allowing for the emergence of outlier cells contributing to population proclivity^25^. On a smaller scale, noise in the expression of individual genes has also been found to be equally important; in *B. subtilis*, controlling transcriptional and translational noise of *comK* was associated with vegetative- and competent-state transitions^27^. Likewise, in cancer, noise can play a significant role, as evidenced by the increasing expression diversity observed in late-stage tumors and their association with cancer outcomes^18,23,24^.

Gene expression noise level affect cell-state transition, in a way similar to the effect of temperature in state transition in inorganic matter. In addition to such ‘standard’ noise following continuous distribution, a ‘discrete’ noise coming from toggle genes^5^ is at play. These genes exhibit a “switch-like” behavior, being “OFF” in one sample (replicate or condition) and “ON” in another, leading to significant weighted noise across samples. This phenomenon has been observed across a wide range of organisms, from unicellular to human mammalian cells, and appears to be consistent regardless of the RNA extraction method employed (Table S1, S2). Of particular interest, toggle genes show a higher incidence in cancer and cell proliferation data, where they contribute significantly to transcriptome-wide noise^5^. Moreover, our observations indicate a greater prevalence of toggle genes in tumor samples compared to their healthy counterparts (Figure 1C). In various cancers, including but not limited to prostate, lung, and breast cancer, similar molecular switches have been observed that not only contribute to drug resistance but also provide the molecular plasticity required for proliferation, metastasis, and uncontrolled growth^28–32^. Thus, further investigation of switch-like or toggle genes in cancer, especially during periods of proliferation, is crucial for understanding the role of gene expression variability in cellular plasticity.

Thus, in this study, we aim to expand the current understanding of CLL proliferation in the context of transcriptomic plasticity by specifically investigating the influence of toggle genes alongside temporal differential gene (DE) expression analyses. We expand the definition of toggle genes to include comparisons between samples of the same condition, capturing variability in gene expression across distinct samples without requiring replicates. To achieve this, we made use of CLL transcriptomic data from several studies (Table S3), with an increased focus on temporal transcriptomic data from a recent study conducted by *Schleiss et.al* ^33^ that investigated the proliferative signature of CLL patient cells by segregating tumor cells into proliferative cells (PC), and non-proliferative cells (NPC)^33,34^. By leveraging advanced data analytics techniques—ranging from correlation, noise analysis, dimensionality reduction and gene enrichment—our objective is to elucidate the complex interplay, and the role played by toggle genes and differentially expressed genes in CLL proliferation.

## Results

### CLL Transcriptome data

For all considered CLL datasets (Table S3,^33,35,36^), we first performed gene expression filtering using statistical distribution fitting and threshold-based filtering (Figure S1, Methods)^16,37^. From the whole transcriptome, this process removed very low and technically noisy genes, leaving only robust gene expressions for further analyses (Table S3). The same was done for the CLL proliferating cells (PC) and non-proliferating cells (NPC) at 9 time points after B cell receptor (BCR) stimulation (*n* = 0, 1, 1.5, 3.5, 6.5, 12, 24, 48, 96h, GSE130385).

### The presence of toggle genes in CLL data

Toggle genes were identified in all three CLL datasets by comparing gene expressions between distinct patient samples exposed to the same disease state. These were termed as toggle genes from same-condition samples, that is, genes with zero expression in one sample and positive expression above a noise threshold in another (Figure 2A). The noise threshold was derived using statistical distribution fitting analysis (Methods), to ensure that the identified toggle genes reflect genuine biological variability rather than technical noise.

**Figure 2.**
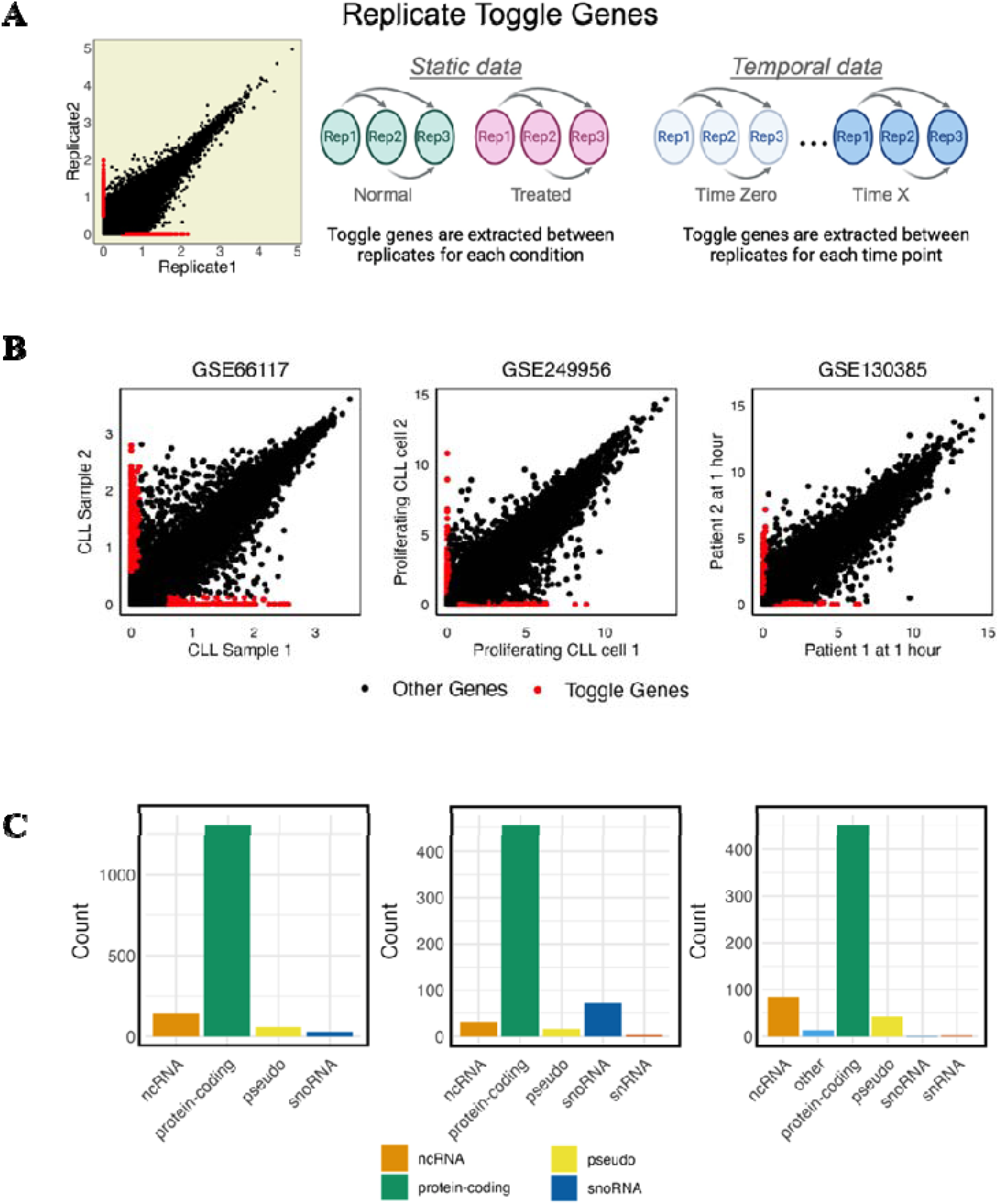
(A) Illustrative schematic of the extraction of toggle genes within the larger transcriptome. (B) Presence of toggle genes (red) within the transcriptome-wide scatter in three investigated CLL datasets: GSE66117, GSE249956, GSE130385. (C) Biotype distribution of toggle genes in three CLL datasets (Table S3), with the prevalent biotype being protein-coding genes.

In the transcriptome-wide scatterplots (Figure 2B), toggle genes (red) are distributed along the x- and y-axes in all datasets. Biotype analysis revealed that the majority of these genes are protein-coding, irrespective of the RNA extraction method used (Table S3), while a smaller subset consists of non-coding genes. The consistent identification of toggle genes in all datasets, combined with their predominance as protein-coding genes, highlights the inherent randomness and instability within CLL transcriptomes.

The concept of ‘randomness’ in toggle genes exhibits a unique characteristic. Typically, randomness is associated with statistical distributions such as uniform, normal, or, more commonly in biological systems, log-normal continuous distributions. However, toggle genes introduce a different form of randomness: toggling is inherently a discrete binary process at the single-cell level. When this behavior extends to the population level in the form of unbalanced toggles, it leads to a pronounced symmetry breaking within the population, ultimately driving the system in a specific direction^38^.We will explore this concept further in the following discussion.

### Tracking the temporal global, DE and toggle genes response

As cell proliferation is a dynamic process, we next investigated the behavior of toggle genes in CLL proliferation using the PC and NPC dataset. DESeq2 analysis identified 9,148 temporal DE genes between time points *t_0_*and *t_n_*, applying a two-fold change and a *p-*value below 0.05 (Table 1). While not unexpected, this substantial gene set, representing 71% of the filtered transcriptome, suggested extensive involvement of DE genes in cell proliferation. To facilitate comparison with the smaller toggle gene set (1,704 genes), the threshold was increased to a four-fold change, reducing the DE gene set to 2,713 genes (Table 1). This stricter threshold helps exclude genetic elements that merely follow the system’s general dynamics due to inter-gene correlations^39^, without being directly involved in the phenomenon under investigation.

**Table 1.** Number of extracted toggle genes and DE genes.

| Gene Set | Number of Toggle Genes |  |  |  |  |  |  |  |  |  |
| --- | --- | --- | --- | --- | --- | --- | --- | --- | --- | --- |
|  | Total: | 0h | 1h | 1h30mi<br>n | 3h30mi<br>n | 6h30mi<br>n | 12h | 24h | 48h | 96h |
| Same-condition<br>Toggle<br>Genes | 1704 | 621 | 610 | 554 | 578 | 545 | 301 | 383 | 285 | 309 |
| Temporal<br>DE<br>Genes,<br>FC>2 | 9148 | NA | 689 | 2580 | 4547 | 3986 | 3656 | 3643 | 5007 | 4600 |
| Temporal<br>DE<br>Genes,<br>FC>4 | 2713 | NA | 179 | 515 | 1171 | 1177 | 1077 | 1184 | 1422 | 1319 |

Subsequent temporal Pearson and Spearman correlation analyses of the transcriptome, toggle genes, and temporal DE genes revealed a rapid decline in correlation between 3.5 and 6.5 hours, followed by stabilization (Figure 3A-C, Pearson, Figure S2, Spearman). Both PC and NPC groups exhibited similar effects, particularly during the critical first four time points. Toggle genes, despite deriving through comparison between same time point and same samples showed dynamic responses similar to global and temporal DE genes. Notably, 604 overlapping genes between toggle and temporal DE genes displayed the most significant correlation drop between 12 and 24 hours, nearly reaching zero, before partial recovery (Fig. 3D). After removing these overlapping genes, the unique toggle genes demonstrated a more pronounced response than the unique temporal DE genes (Fig. 3E-F). This indicates that toggle genes, on top of DE genes, contribute significantly to the temporal PC and NPC responses.

**Fig. 3.**
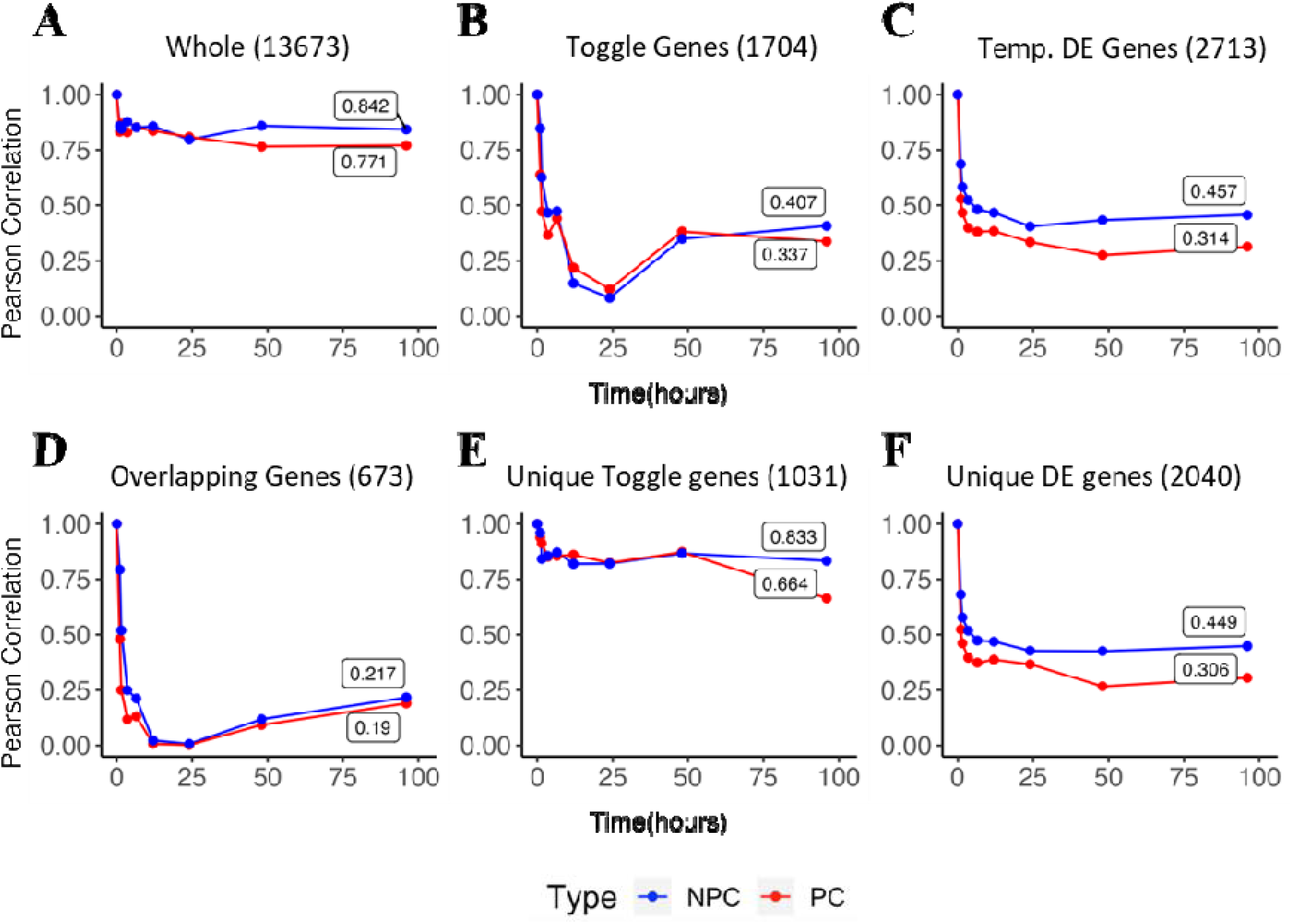
Average autocorrelation of PC and NPC cells across time. (A) Pearson autocorrelation for the whol transcriptome (13K genes). (B) Pearson autocorrelation for extracted toggle genes (1.7K genes). (C) Pearson autocorrelation for high fold-change (4FC) differentially expressed temporal genes (3K genes). (D) Pearson autocorrelation for the overlapping genes between DEG and Toggle genes. (E) Pearson autocorrelation for unique toggle genes. (F) Pearson autocorrelation for Unique DEG.

Overall, these results suggest that the pronounced changes in correlation and auto-correlation observed for toggle genes and DE genes reflect the proliferative processes occurring within CLL cells. Both gene sets exhibit significantly larger responses compared to the rest of the transcriptome, with their intersection capturing some of the most dynamically responsive genes in both PC and NPC groups.

### Toggle genes possess the highest gene expression noise

Gene expression noise, measured as the squared coefficient of variation (CV²), was evaluated for the whole transcriptome and specific gene sets, including DE, toggle, overlapping, and random subsets (Methods). Two types of noise were assessed: (1) **between-sample noise**, which quantifies variability among samples of the same condition (e.g., PC or NPC) at the same time point, and (2) **temporal noise**, which evaluates variability across time by comparing each time point to the baseline transcriptome at *t_0_*(Fig. 4, Figure S3).

**Fig. 4.**
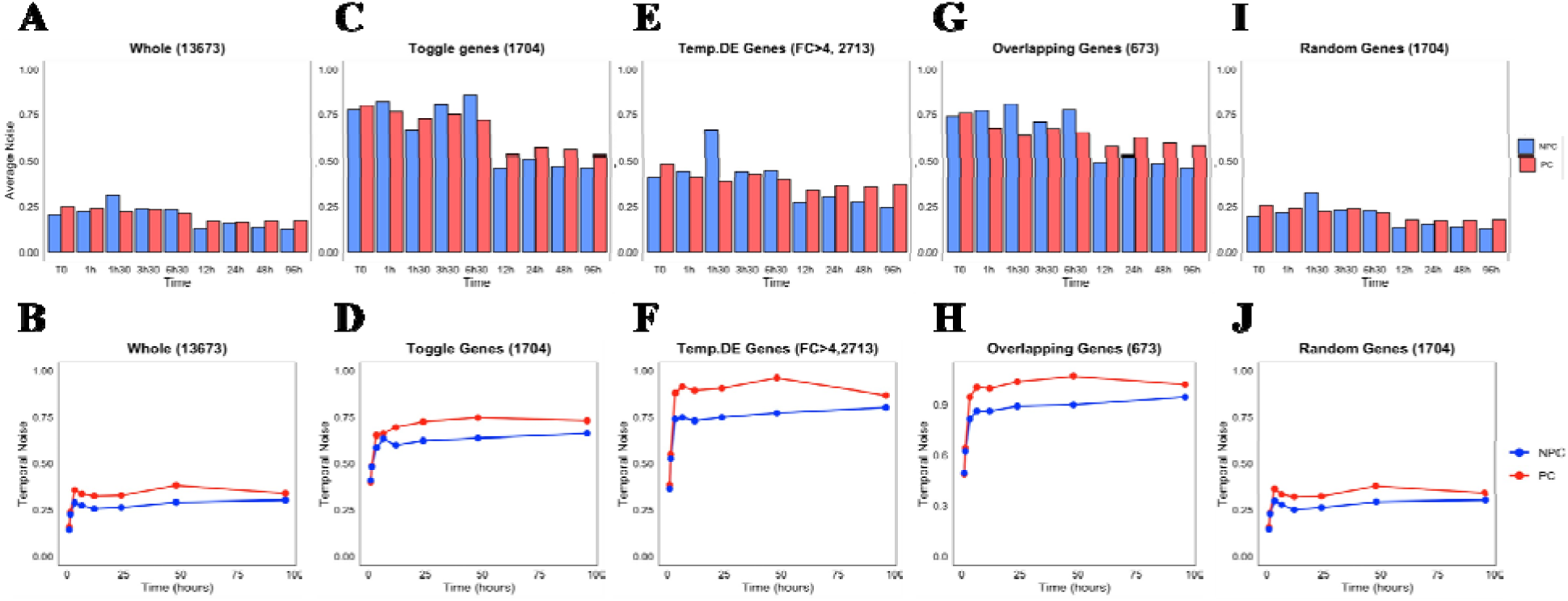
Noise changes in time for PC (red) and NPC (blue) samples. (A) Transcriptome-wide average noise changes in time between same-condition samples. (B) Average noise changes relative to time zero for the whol transcriptome. (C) Average noise changes in time between samples for toggle genes. (D) Average noise changes relative to time zero for toggle genes. (E) Average noise changes in time between samples for high FC DE genes. (F) Average noise changes relative to time zero for 4FC DE genes. (G) Average noise changes in time between samples for overlapping DE and toggle genes. (H) Average noise changes relative to time zero for overlapping DE and toggle genes. (I) Average noise changes in time between samples for random subsets of 1704 genes. (J) Average noise changes relative to time zero for random subsets of 1704 genes.

For between-sample noise, toggle genes exhibited the highest variability levels, followed by DE and overlapping genes, in both PC and NPC groups. Noise levels peaked at 6.5 hours post-stimulation across all gene sets, suggesting increased variability among same-condition samples at this time point. This heightened variability reflects greater heterogeneity within the population, which stabilized at later time points (Fig. 4a, c, e, g, i, S3).

Temporal noise analysis showed that DE genes exhibited slightly higher levels than toggle genes, with both sets displaying significantly greater noise compared to the whole transcriptome or random subsets (Fig. 4b, d, f, h, j, S3). Notably, overlapping genes, despite representing only a small fraction of the other gene sets, exhibited the greatest temporal changes, highlighting their substantial contribution to transcriptome-wide noise and their distinct dynamic behavior over time. Notably, the increased temporal variability of toggle genes is intrinsically linked to their bi-stable nature, causing them to oscillate between two extremes. This characteristic makes them natural ‘noise amplifiers,’ particularly when an imbalance occurs in their oscillation between ON and OFF states^38^.

To go deeper into gene expression variability, Shannon entropy was analyzed for the same gene sets (Methods)^40^. The whole transcriptome and random subsets exhibited lower entropy levels with no discernible temporal patterns (Figure S4). By contrast, toggle and DE genes showed higher entropy levels but displayed distinct temporal trends. Toggle genes demonstrated a steady decline in entropy, reaching their lowest levels at 24 hours, while DE genes showed a gradual increase, stabilizing by 24 hours. Overlapping genes exhibited the most pronounced downward trend, indicating a stronger dynamic response than either toggle or DE genes. Random subsets showed a pattern similar to the whole transcriptome but with slightly higher entropy due to their smaller size. These results suggest that toggle and DE genes contribute significantly to expression variability within samples, with overlapping genes displaying unique dynamic behavior.

Lastly, we analyzed temporal toggle genes, defined as genes toggling in expression between time points (*t_0_* and *t_n_*). A total of 2,561 temporal toggle genes were identified. However, noise and autocorrelation analyses revealed weaker responses for temporal toggle genes compared to same-condition toggle genes, likely due to differences in the size and composition of the gene sets (Figures S3–S4). Despite this, the analysis of temporal toggle genes provides additional insights into transcriptomic variability over time and emphasizes the complexity of gene expression dynamics in CLL.

### Gene Enrichment analyses of toggle and DE genes for PC and NPC

Now that we have shown both toggle and DE genes are important for shaping temporal dynamics and variability in CLL, to understand their biological functions, the *Reactome* pathway enrichment analysis was conducted. Toggle genes were enriched in key processes such as lymphoid cell communication and RHO GTPases, while DE genes were associated with immune-related pathways, including interleukin signaling and TNF-related processes (Fig. 5A - toggle genes, 5B - DEGs, 6C - overlapping genes). Notably, overlapping genes, which shared characteristics of both toggle and DE genes, were particularly enriched in chemokine receptor processes, interleukin signaling, and lymphoid immunoregulatory interactions.

**Fig. 5.**
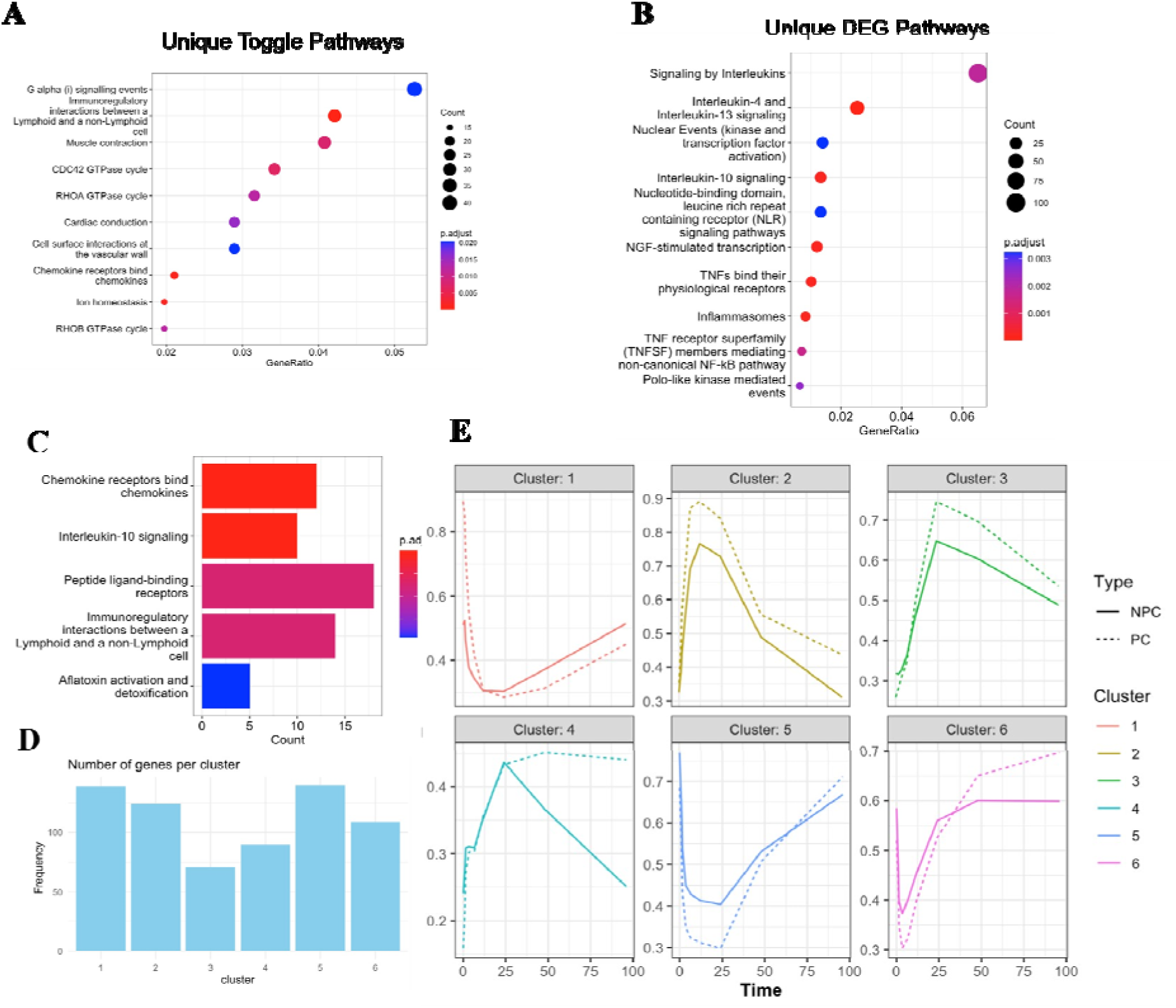
Functional enrichment analysis of DE genes and toggle genes and the overlap between the different sets. (A) Enriched Reactome pathways for unique toggle genes. (B) Enriched Reactome pathways in temporal DE genes. (C) Enriched Reactome pathways for overlapping DEG and toggle genes. (D) Average temporal signatures of overlapping gene clusters based on hierarchical clustering.

Given that the experimental setup involved cell treatment with chemokines and interleukins to stimulate survival and proliferation, the enrichment of these processes among toggle and overlapping genes serves as a proof of principle, underscoring their biological significance. This alignment between the observed enrichment and the experimental conditions reinforces the importance of toggle genes in the cellular responses studied.

Further analysis of overlapping genes identified six clusters ranging from 70 to over 100 genes, each with distinct temporal expression profiles (Figure 5D). Sharp early responses were observed in interleukin signaling and chemokine-related processes, particularly in clusters 1 and 2 (Figure S5 A, B). Additionally, cell cycle checkpoint processes exhibited a delayed response, peaking at 24 hours before declining, consistent with the major transcriptomic changes noted in earlier analyses (S5).

The following are top 10 toggle genes based on their squared of variation (CV): SOX2, NCS1, ALPP, GPR34, EEPD1, SPNS2, CYP2C18, SIX3, F2RL2, RPRML. Notably, SOX2, NCS1 and SPNS2 stand out for their potential involvement in the proliferation of CLL cells. SOX2, a transcription factor that is necessary for maintaining stem cell properties, has been shown to contribute to the self-renewal and tumorigenic potential of leukemia stem cells^41^. NCS1 (neuronal calcium sensor 1) encodes a protein that regulates calcium signaling, which has been found to be essential for immune cell function and activation, with its dysregulation potentially driving leukemogenesis^42^. Lastly, SPNS2, involved in transporting sphingosine-1-phosphate (S1P), affects leukemic cell migration and survival, which are both essential for CLL cells in the lymph node microenvironment^43^. The genes identified, such as SOX2, NCS1, and SPNS2, influence critical processes like cell signaling, migration, and self-renewal in CLL cells, all of which contribute to CLL progression and provide potential targets for future therapeutic strategies.

In summary, the enrichment analysis demonstrates that toggle genes, especially those overlapping with DE genes, are involved in critical biological processes related to immune function, cell cycle regulation, and differentiation, aligning closely with the experimental conditions designed to activate these pathways.

## Discussion

The study of cancer presents significant challenges not only because of the disease’s inherent complexity and aggressiveness but also due to its heterogeneous nature, including cellular plasticity, compounded by a limited understanding of transitions between cancer states^8,10^. Cellular plasticity and state transitions are thought to be influenced by transcriptomic instability, which has been previously linked to tumor progression and treatment resistance^9,12^. As observed in previous studies, the transcriptomes of cancer cells are often unstable and display unique expression deviations^44–46^. This underscores the need for approaches that capture transcriptomic variability, including noise, which has been shown to play a role in shaping cell states and tipping cellular trajectories^15,18,21,25,27^.

Molecular “switch-like” behaviors, characterized by flexibility and plasticity, have been shown to contribute to adaptive and evasive behaviors in cancer cells^29,31,32^. Toggle genes, which exhibit binary “ON/OFF” expression patterns, represent a specific instance of this phenomenon. Our findings showed an increased incidence of toggle genes in cancer samples compared to healthy or adjacent tissues from the same individuals. This observation highlights the variability within cancer transcriptomes, which may reflect broader processes like proliferation or immune modulation. On a more general perspective, the higher proportion of toggle genes in cancer is consistent with the ‘noise amplifier’ role allowing cancer cells to explore a wider phase space exploration than healthy cells. This noise amplification has very important consequences in terms of therapy resistance and recurrence of cancer^8^

Toggle, or bi-stable, genes have been shown to play a key role in regulating complex biological systems in various instances. Research on these genes began with the well-known example of ON/OFF regulation: the alternation between the lytic and lysogenic phases of phage lambda^47,48^. It is also known that many endogenous retrovirus (ERV) sequences exhibit a bi-stable (yes/no) activation behavior, inherited from their viral origins, which could potentially be leveraged in cancer therapy^49^.The frequency of ERVs positively correlates with evolutionary complexity and varies significantly between cell lines.^50,51^

By focusing on the temporal transcriptomic dynamics of CLL cells following BCR stimulation—a key driver of proliferation in this disease—we sought to investigate how toggle genes and transcriptomic noise contribute to variability during the proliferative response. Rather than implying causality, we aimed to show that these transcriptomic features align and correlate with the instabilities observed during CLL proliferations.

We identified 1,704 toggle genes and 2,713 DE genes with a significant temporal response (above 4-fold change). Auto-correlation analysis revealed a sharp decline in transcriptome correlation between 3.5 and 6.5 hours post-stimulation, coinciding with early proliferation events. This pattern of variability, particularly in PCs, suggests that transcriptomic instability accompanies the proliferative process. A subset of toggle genes overlapped with DE genes, showing the largest temporal shifts, while unique toggle genes displayed variability across same-condition samples. This distinction was further supported by dimensionality reduction, noise, and entropy analyses, which revealed that overlapping genes exhibit characteristics of both toggle and DE genes. These findings reinforce the idea that transcriptomic instability underlies the dynamic responses observed during CLL proliferation.

The enrichment analysis provided additional insight into the biological relevance of toggle and overlapping genes. The enrichment of toggle-genes involved in G-alpha signaling, muscle contraction, and cardiac conduction could be considered as largely unexpected, while the enrichment of chemokine and interleukin signaling, aligns with the experimental conditions designed to promote survival and proliferation. In this respect, it is worth noting that cytoskeleton remodeling (driven by the same genes linked to muscel contraction) is since long time recognized as a crucial player in cancer^52^ while being in the same time an obliged step in cell division. Similar considerations hold for cardiac conduction genes^53^ and G-alpha signaling^54^. The presence of differentially enriched pathways validates the notion that toggle-genes observed variability reflects biologically meaningful responses rather than pure random noise. Furthermore, the enrichment of RHO-GTPase signaling suggests potential novel mechanisms underlying cancer proliferation, offering new directions for investigation.

Interestingly, the overlapping genes represent a subset of the transcriptome that bridges temporal responsiveness and variability across samples. This dual role highlights their importance in both proliferation and plasticity. For instance, processes like chemokine signaling, which are well-established in CLL, were also enriched in toggle genes, indicating their potential contribution to both immune modulation and cellular heterogeneity. This supports the hypothesis that toggle genes reflect disturbances within important processes as evidenced by their transcriptomic expression, that can contribute to variations in disease progression.

Finally, our findings on RHO-GTPases underscore their significance in cancer dynamics^55^. Their consistent temporal expression patterns, coupled with differences between PC and NPC groups, suggest they play a regulatory role in tumor initiation and progression. These genes, identified as toggle genes in this study, may serve as key regulators of cellular behaviors essential for cancer development, making them potential therapeutic targets in CLL.

Overall, our study highlights the role of transcriptomic instability as a feature of cancer proliferation. Toggle genes, particularly those overlapping with DE genes, provide evidence of this instability, reflecting both temporal changes and population-level variability. By identifying the dynamic interplay between noise, gene expression dynamics, and cellular behavior, this study deepens our understanding of CLL’s proliferative signature and its complex molecular underpinnings from a system dynamics viewpoint. Future work should further explore these transcriptomic features to uncover their impact on disease progression and actual treatment outcomes, with the aim of developing more targeted novel therapeutics.

## Methods

### Pre-processing

For the time series data (*GSE130385*^33^), we first removed genes with constant zero expression in all samples (24,477), and performed trimmed mean of M values (TMM) normalization^56^ on the remaining gene counts. Gene expression distribution fitting was then performed using *fitdistplus*^57^ , and *mass*^58^, for several distribution types: log-normal, log-logistic, Pareto, Burr and Weibull. Lastly, an expression cut-off was identified and used to filter for genes with expression above the cut-off in at least one sample, with the final number of genes being 13,673.

### Toggle gene extraction

Same-condition sample toggle genes were identified and extracted as defined by Giuliani, et al^28^:

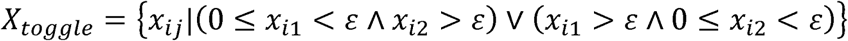

where, *x_ij_* represents the expression vector of the *i*-th gene for two samples *j*=1,2 of the same condition. The parameter *ε* denotes the minimum expression threshold determined from statistical distribution fitting step above.

Similarly, temporal toggle genes were extracted using the same criteria across different time (*n*) points of the same condition: *j*= 0, *n^th^* time point.

For each condition with three biological samples, toggle genes are identified pairwise, meaning that a gene may toggle between any two samples within the condition without requiring toggling across all sample pairs. Similarly, temporal toggle genes were extracted using the same criteria but applied across different time points of the same condition: *j* = *t_0_t_1_*, *t_0_t_2_*, ….,*t_0_t_n_*, where *t_0_t_n_* represents comparison between *t_0_*and *t_n_* time points, comparing all time points with final time *t_n_*. This approach ensures that toggling behavior is evaluated consistently across both same-condition and temporal contexts.

### DE gene extraction

Temporal DE genes were extracted using *DESeq2*^59^, using a fold-change of 2 and 4 as indicated in maintext. DE analysis was performed between the initial time points (*t_0_*) and the *n*-th time points (*t_n_*), where *n* > 0, for both PC and NPC conditions. Only genes that passed a threshold of *p-*value < 0.05 were retained.

### Correlation

Autocorrelation refers to correlation changes with respect to *t_0_*and is computed by calculating the correlation between *t_0_* and *t_n_* respectively. Two auto-correlation metrics were deployed in this analysis: Pearson correlation and Spearman correlation.

### Pearson

Pearson correlation between two vectors can be calculated as:

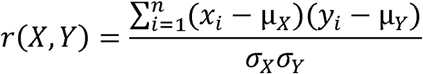

where μ*_X_* and μ*_Y_* are the mean values for vectors ***X*** and ***Y***, and similarly *σ_X_* and *σ_Y_* represent the standard deviations. In the case of autocorrelation, ***X*** always refers to the initial time point, and ***Y*** to each subsequent time point.

### Spearman

Like Pearson correlation, Spearman rank correlation between ***X*** and ***Y*** is defined as:

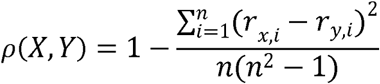

Where *r_x,i_* and *r_y_*_,*i*_ represent the ranks of the *i-*th observation in the initial time point and the considered time point.

### Noise

Noise between any two samples was computed using:

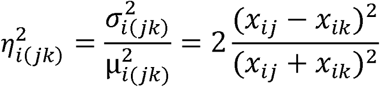

where *X_ij_* and *X_ik_* are the values of a gene in *j-*th and *k-*th samples. Average noise is calculated by averaging the summed noise values of all genes between all pairs considered giving a final noise formula of:

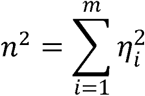

For temporal noise, the calculation was performed for each time point with respect to *t_0_,* and for sample noise, the calculation was performed between all samples of any given sample condition.

### Hierarchical Clustering

Hierarchical clustering for toggle genes and DEG was performed using the *stats* package in R, where first a distance matrix between the samples was computed for each corresponding gene set. Next, Ward clustering^60^ method was applied to group genes with similar temporal expression patterns. For each identified cluster, the mean TMM expression across all timepoints was plotted to visualize temporal expression patterns for both PC and NPC.

### GO and Network Analysis

For GO analysis, several analytic tools were performed. Gene enrichment analysis for Biological Processes was performed using *clusterProfiler*^61^ in R, using a threshold of *p-value* < 0.05. Next, GO networks were generated in Cytoscape using ClueGO^62^, with specificity chosen as global, and a significance threshold below 0.05. Lastly, *Reactome*^63^ pathway analysis was performed to gain further understanding of the enriched pathways within the temporal proliferative signature with a similar threshold of *p-*value <0.05.

## Data Availability

The codes for the analysis used in this manuscript are available from the authors upon request. The CLL transcriptomic data used in this manuscript can be found in the GEO database using the accession numbers: GSE66117, GSE249956, GSE130385.

## Author Contributions

Olga Sirbu: Conceptualization; writing – original draft; formal analysis (lead); conceptualization (supporting) Gunjan Agarwal: Writing – reviewing and editing; formal analysis (supporting); conceptualization (supporting) Alessandro Giuliani: Conceptualization (supporting) ; writing - reviewing and editing (supporting) . Kumar Selvarajoo: Conceptualization (lead); writing - reviewing and editing (lead).

