## Supplementary Figure for "Understanding the Role of Toggle Genes in Chronic Lymphocytic Leukemia Proliferation"

***Table S1. RNA extraction method for studies used***

| **Species/cell type** | **GEO accession** | **RNA Extraction Method** |
|---|---|---|
| *Saccharomyces cerevisiea* | GSE85595 | Poly(A) Selection |
| *Escherichia coli* | GSE71562 | rRNA Depletion |
| *Drosophila melanogaster* | GSE167197 | Poly(A) Selection |
| *Bacillus subtilis* | GSE141305 | rRNA Depletion |
| *Canis lupus familiaris* | GSE162281 | Poly(A) Selection |
| *Arabidopsis thaliana* | GSE155365 | Poly(A) Selection |
| *Homo sapien T-regulatory* | GSE94396 | Poly(A) Selection |
| *Mus musculus macrophage* | GSE160640 | Poly(A) Selection |
| *Danio rerioembryo* | GSE147112 | Poly(A) Selection |
| *Yarrowia lipolitica* | GSE151659 | Poly(A) Selection |
| *Gallus gallus* | GSE135272 | rRNA Depletion |
| *Sus scrofa* | GSE159583 | Poly(A) Selection |
| *Caenorhabditis elegans* | GSE149862 | Poly(A) Selection |
| *Bos taurus* - cloned embryo | GSE123705 | Poly(A) Selection |
| *Homo sapien* - Kaposi sarcoma | GSE100684 | Poly(A) Selection |
| *Mus musculus* - lung cancer | GSE158502 | Poly(A) Selection |
| *Macaca mulatta* - brain | GSE120901 | rRNA Depletion |
| *Homo sapien* - myeloma | GSE129801 | Poly(A) Selection |
| *Homo sapien* - lymphoma | GSE148656 | Poly(A) Selection |
| *Mus musculus* - thyroid cancer | GSE162795 | Poly(A) Selection |
| *Macaca mulatta* - blastocyst | GSE162817 | Poly(A) Selection |
| *Homo sapien* - oocyte | GSE155489 | Poly(A) Selection |
| *Homo sapien* - embryonic cardiomyocyte | GSE137255 | Poly(A) Selection |

***Table S2. Datasets used for figure 1C, including accession number, cancer type, sequencing type, as well as RNA extraction protocol.***

| Accession Number | Cancer Type | Sequencing Type | RNA Extraction Method |
| --- | --- | --- | --- |
| GSE183904 | Gastric | Single cell | polyA |
| GSE126209 | Osteosarcoma | Bulk | polyA |
| GSE112705 | Liver | Bulk | polyA |
| GSE129801 | Multiple myeloma | Bulk | RNA depletion |
| GSE148656 | Lymphoma | Bulk | polyA |
| GSE100684 | Sarcoma | Bulk | polyA |
| GSE184616 | Skin | Bulk | RNA depletion |
| GSE190688 | Ovarian | Bulk | polyA |

***Table S3. Number of genes before and after distribution-fitting based filtering for the selected CLL studies***

| **GEO Accession Number** | GSE66117 | GSE249956 | GSE130385 |
|---|---|---|---|
| **No of samples** | 14 (11 CLL patients, 3 healthy donors) | 40 samples (Proliferative and resting fraction cells from CLL patients; 3 samples per condition) | 54 samples (2 conditions: PC and NPC; 3 samples per condition, across 9 time points) |
| **Genes before filtering** | 22 249 | 60 624 | 27 253 |
| **Genes after filtering** | 13670 | 25369 | 13 673 |
| **Filtering Threshold** | 0.5 | 7 | 5 |
| **Extraction Method** | poly-A purification | poly-A purification | rRNA depletion |
| **Study Setup** | CLL transcriptome comparison between patient and healthy donor samples | Longitudinal analysis of PF and RF cells in CLL patients treated with ibrutinib | Time-series profiling of CLL cells post BCR stimulation across nine time points for proliferative (PC) and non-proliferative (NPC) cells |

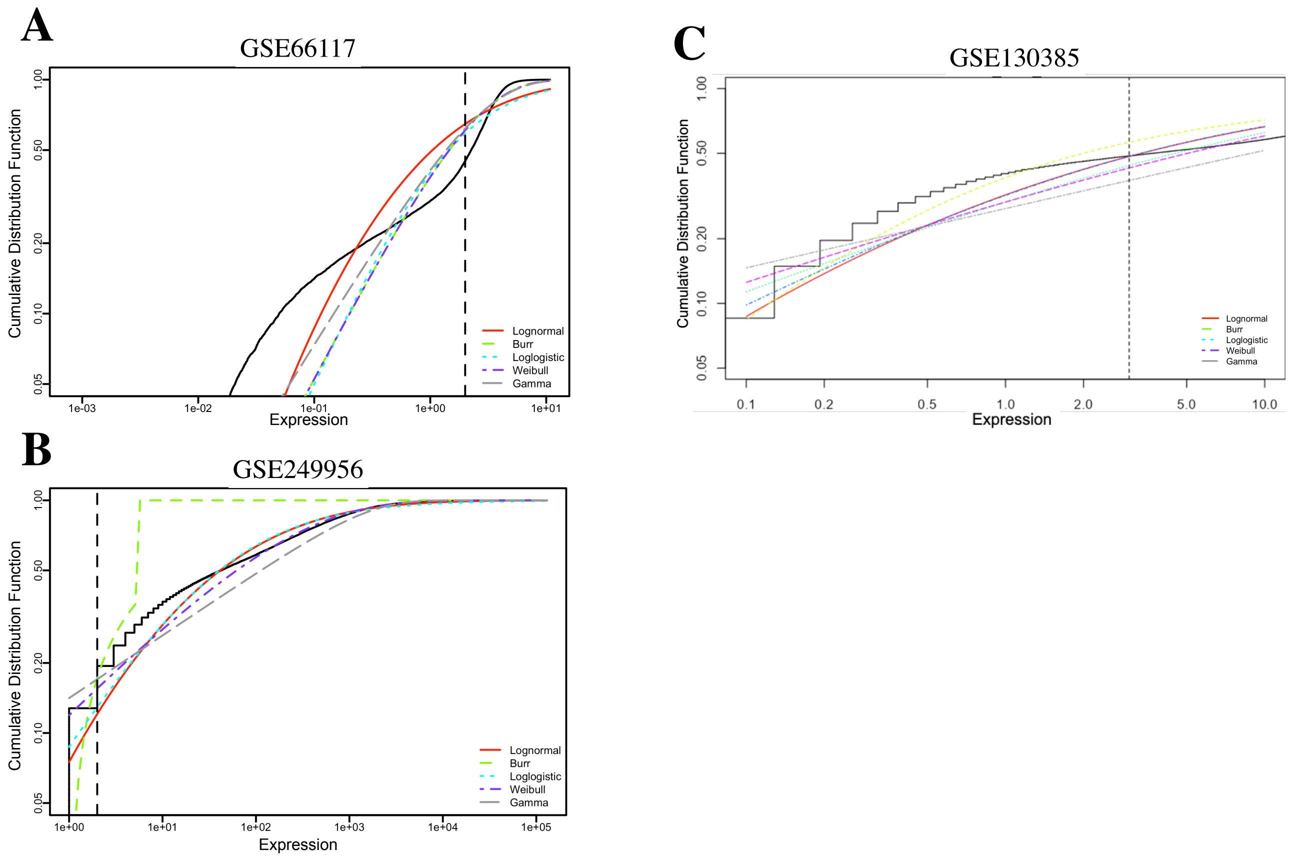


***Fig. S1.*** Cumulative Distribution Functions for expression values of three CLL datasets (A-C), with experimental data in black, lognormal in red, burr in green, log logistic in cyan, Weibull in purple and Gamma in grey. The vertical dotted line represents the intersection the selected threshold for filtering.

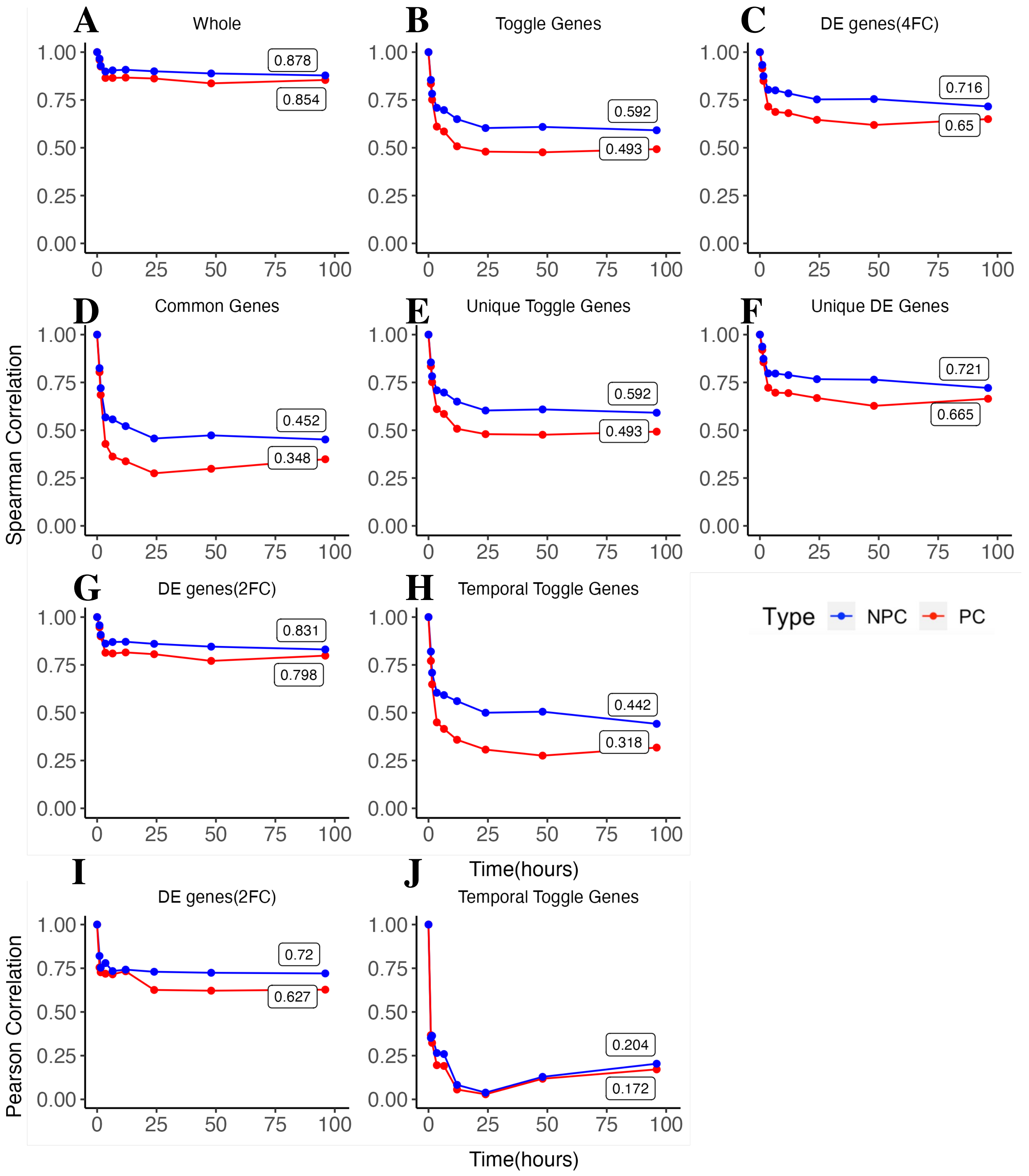


**Fig. *S2*** Average auto-correlation of PC and NPC cells across time. (A) Spearman auto-correlation for the whole transcriptome (13K genes). (B) Spearman auto-correlation for extracted toggle genes (1.7K genes). (C) Spearman auto-correlation for high fold-change (4FC) differentially expressed temporal genes (3K genes). (D) Auto-correlation for the overlapping genes between DEG and Toggle genes. (E) Unique toggle genes. (F) Unique DEG. (G) Spearman auto-correlation for low fold-change (2FC) differentially expressed temporal genes. (H) Spearman auto-correlation for toggle genes. (I) Pearson auto-correlation for low fold-change (2FC) differentially expressed temporal genes. (J) Pearson auto-correlation for toggle genes.

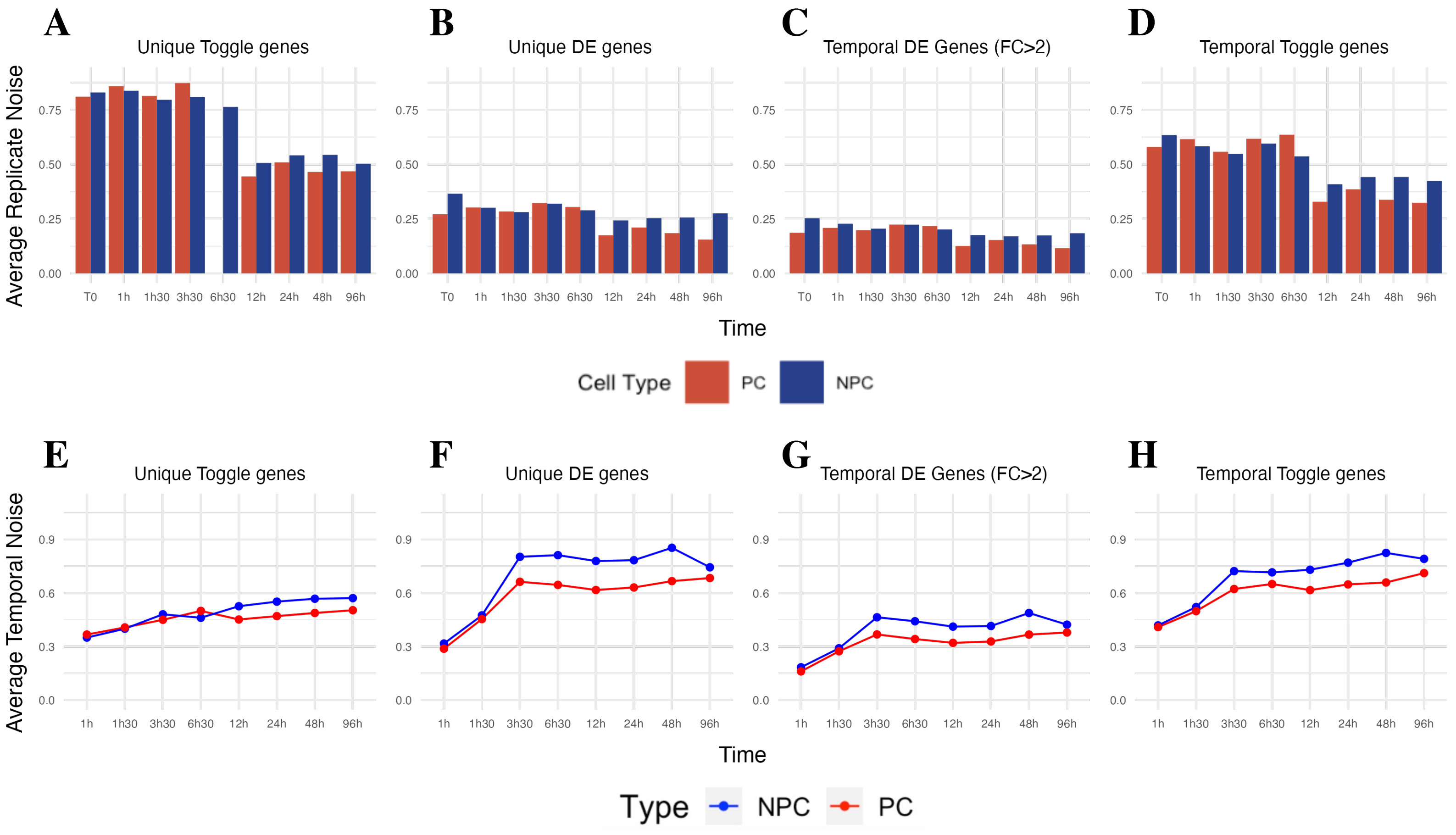


**Fig. S3** Noise changes in time for PC (red) and NPC (blue) samples. (A) Unique toggle genes average noise changes in time between replicates. (B) Unique DE genes average noise changes in time between replicates. (C) Average noise changes in time between replicates for 2FC DE genes. (D) Temporal toggle genes average noise changes in time between replicates. (E) Average noise changes relative to time zero for unique toggle genes. (F) Average noise changes relative to time zero for unique DEG. (G) Average noise changes relative to time zero for temporal 2FC DEG. (H) Average noise changes relative to time zero for temporal toggle genes.

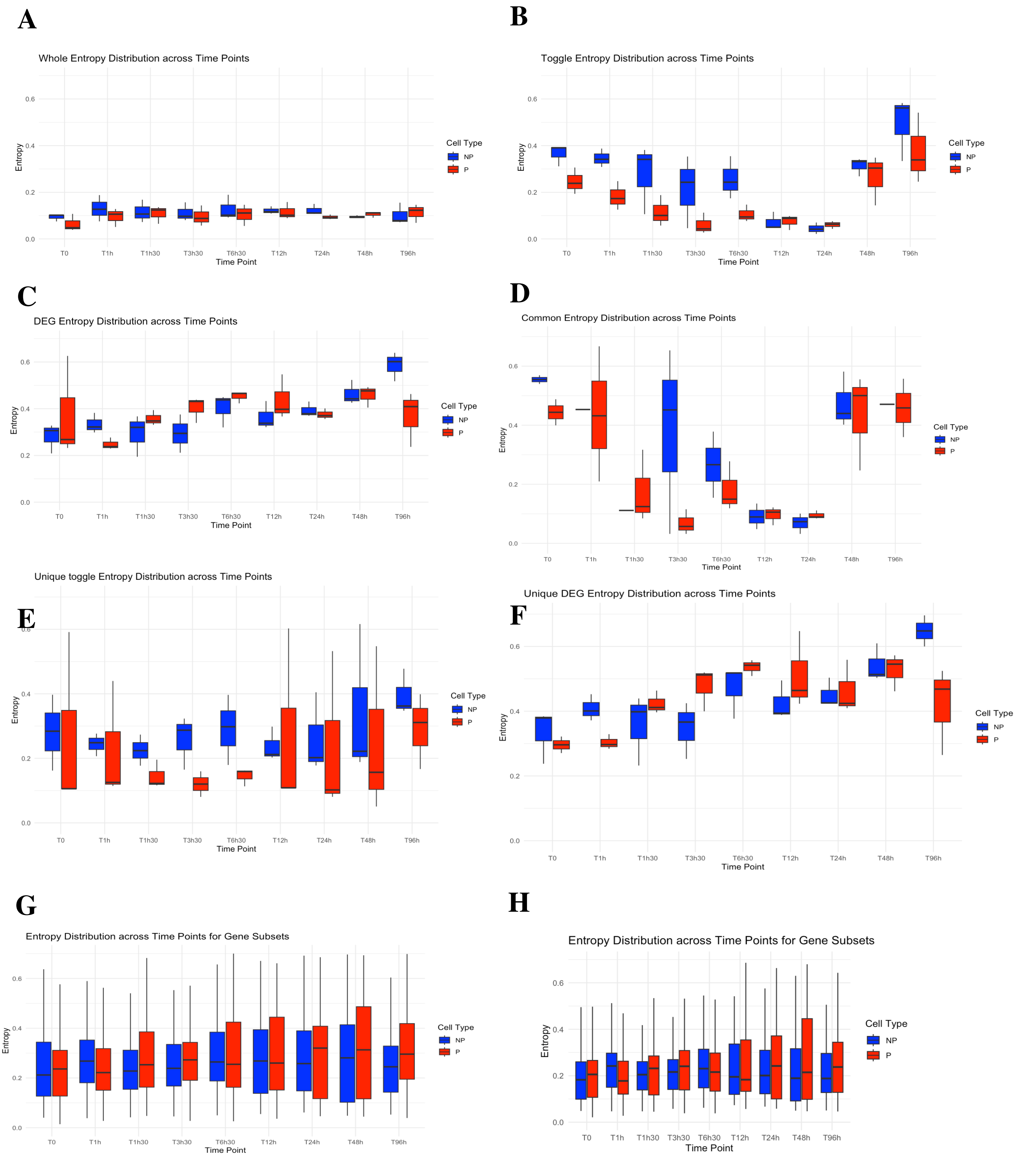


**Fig. S4** Entropy in time for PC (red) and NPC (blue) samples. Average replicate entropy for the (A) whole transcriptome, (B) toggle genes, (C) 4FC DE genes, (D) overlapping DEG and toggle, (E) unique toggle genes, (F) unique DEG, (G) random genes of size n=600, (H) random genes of size n=1704.

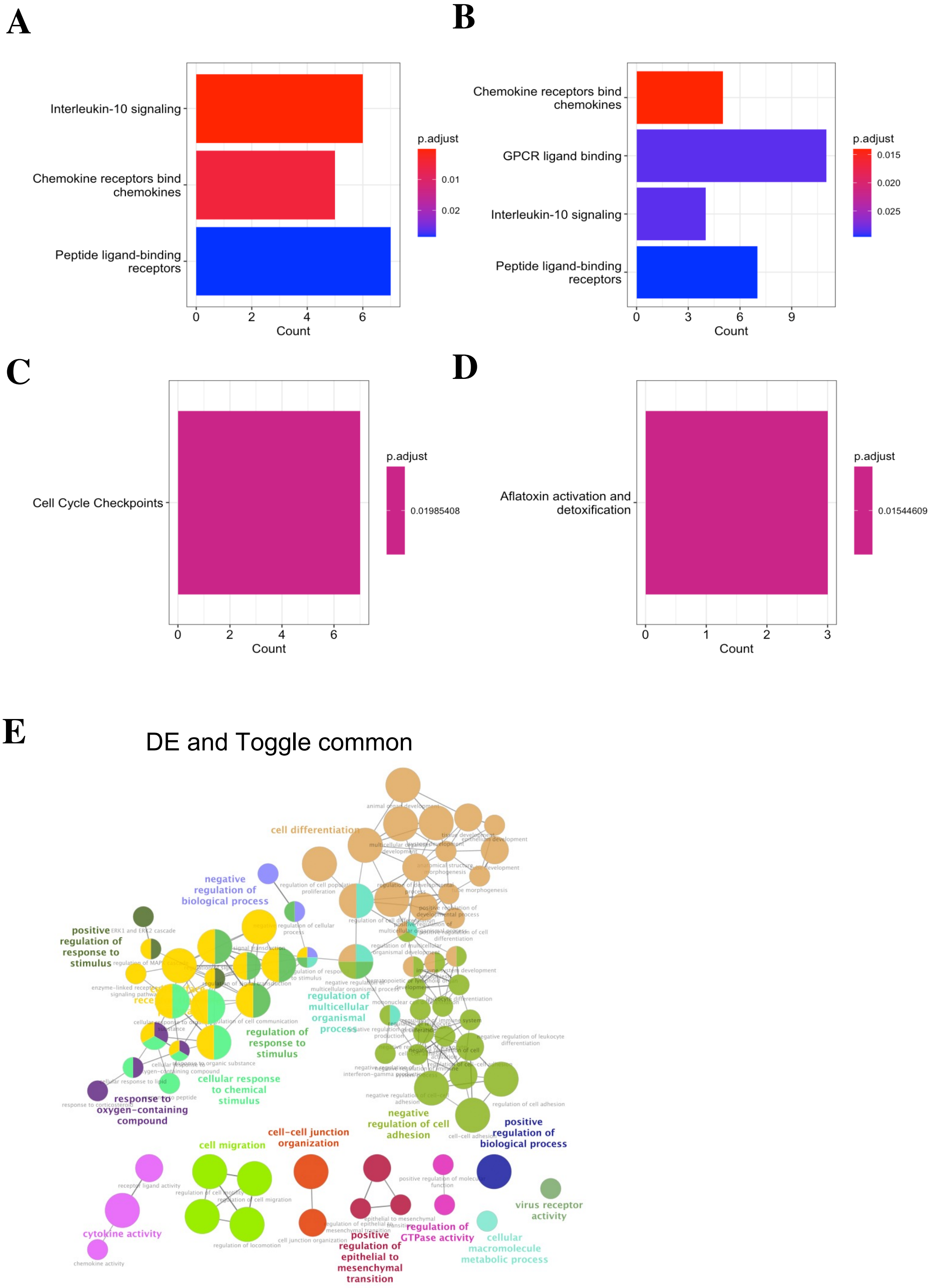


**Fig. S5** Reactome pathway enrichment for (A) cluster 1, (B) cluster 2, (C) cluster 3, (D) cluster 5, from toggle gene based hierarchical clustering. (E) ClueGO based enrichment for overlapping DEG and toggle genes.
